# PESCA: A scalable platform for the development of cell-type-specific viral drivers

**DOI:** 10.1101/570895

**Authors:** Sinisa Hrvatin, Christopher P. Tzeng, M. Aurel Nagy, Hume Stroud, Charalampia Koutsioumpa, Oren F. Wilcox, Elena G. Assad, Jonathan Green, Christopher D. Harvey, Eric C. Griffith, Michael E. Greenberg

## Abstract

Enhancers are the primary DNA regulatory elements that confer cell type specificity of gene expression. Recent studies characterizing individual enhancers have revealed their potential to direct heterologous gene expression in a highly cell-type-specific manner. However, it has not yet been possible to systematically identify and test the function of enhancers for each of the many cell types in an organism. We have developed PESCA, a scalable and generalizable method that leverages ATAC- and single-cell RNA-sequencing protocols, to characterize cell-type-specific enhancers that should enable genetic access and perturbation of gene function across mammalian cell types. Focusing on the highly heterogeneous mammalian cerebral cortex, we apply PESCA to find enhancers and generate viral reagents capable of accessing and manipulating a subset of somatostatin-expressing cortical interneurons with high specificity. This study demonstrates the utility of this platform for developing new cell-type-specific viral reagents, with significant implications for both basic and translational research.

**One sentence summary:** Highly paralleled functional evaluation of enhancer activity in single cells generates new cell-type-specific tools with broad medical and scientific applications.

## Main text

### Introduction

Enhancers are DNA elements that regulate gene expression to produce the unique complement of proteins necessary to establish a specialized function for each cell type in an organism. Large scale efforts to build a definitive catalogue of cell types (*1–4*) based on their gene expression have recently successfully mapped epigenomic regulatory landscapes (*5–7*), enabling a mechanistic understanding of the underlying gene expression that is critical for cell-type-specific development, identity, and unique function. Importantly, characterization of individual enhancers has revealed their potential to direct highly cell-type-specific gene expression in both endogenous and heterologous contexts (*8–11*), making them ideal for developing tools to access, study, and manipulate virtually any mammalian cell type.

Despite recent success in cataloguing the gene expression profiles of distinct cell subpopulations in the nervous system, our limited ability to specifically access these subpopulations hinders the study of their function. For example, the mammalian cerebral cortex is composed of over one hundred cell types, most of which cannot be individually accessed using existing tools. Glutamatergic excitatory neuron cell types propagate electrical signals across neural circuits, whereas GABAergic inhibitory interneuron cell types play an essential role in cortical signal processing by modulating neuronal activity, balancing excitability, and gating information (*12–14*). Although relatively lower in abundance than excitatory neurons, interneurons are highly diverse; for example, somatostatin-expressing cortical interneurons comprise several anatomically, electrophysiologically, and molecularly defined cell types whose dysfunction is associated with neuropsychiatric and neurological disorders (*2, 15, 16*). Given the vast diversity of cell types in the brain, and the inability of our current tools to access most neuronal cell types, enhancer-driven viral reagents have the potential to become the next generation of cell-type-specific transgenic tools enabling facile, inexpensive, cross-species, and targeted observation and functional study of neuronal cell-types and circuits.

Despite the potential of cell-type-specific enhancers to revolutionize neuroscience research, cell-type-restricted gene regulatory elements (GREs) have not yet been systematically identified. Moreover, functional evaluation of candidate GRE-driven viral vector expression across all cell types in the tissue is currently laborious, expensive, and low-throughput, typically relying on the production of individual viral vectors and the assessment of expression across a limited number of cell types by *in situ* hybridization or immunofluorescence. The lack of a generalizable platform for rapid identification and functional testing of cell-type-specific enhancers is therefore a critical bottleneck impeding the generation of new viral reagents required to elucidate the function of each cell type in a complex organism.

To address these issues, we merged the principles of massively parallel reporter assays (MPRA) (*17–22*) with single-cell RNA sequencing (scRNA-seq) (*1*–*4, 23*–*27*), and developed a Paralleled Enhancer Single Cell Assay (PESCA) to identify and functionally assess the *specificity* of hundreds of GREs across the full complement of cell types present in the brain. In the PESCA protocol, the expression of a barcoded pool of AAV vectors harboring GREs is analyzed by single-nucleus RNA sequencing (snRNA-seq) to evaluate the specificity of each constituent GRE across tens of thousands of individual cells in the target tissue, through the use of an orthogonal cell-indexed system of transcript barcoding (**Fig. 1a**).

**Figure 1.**
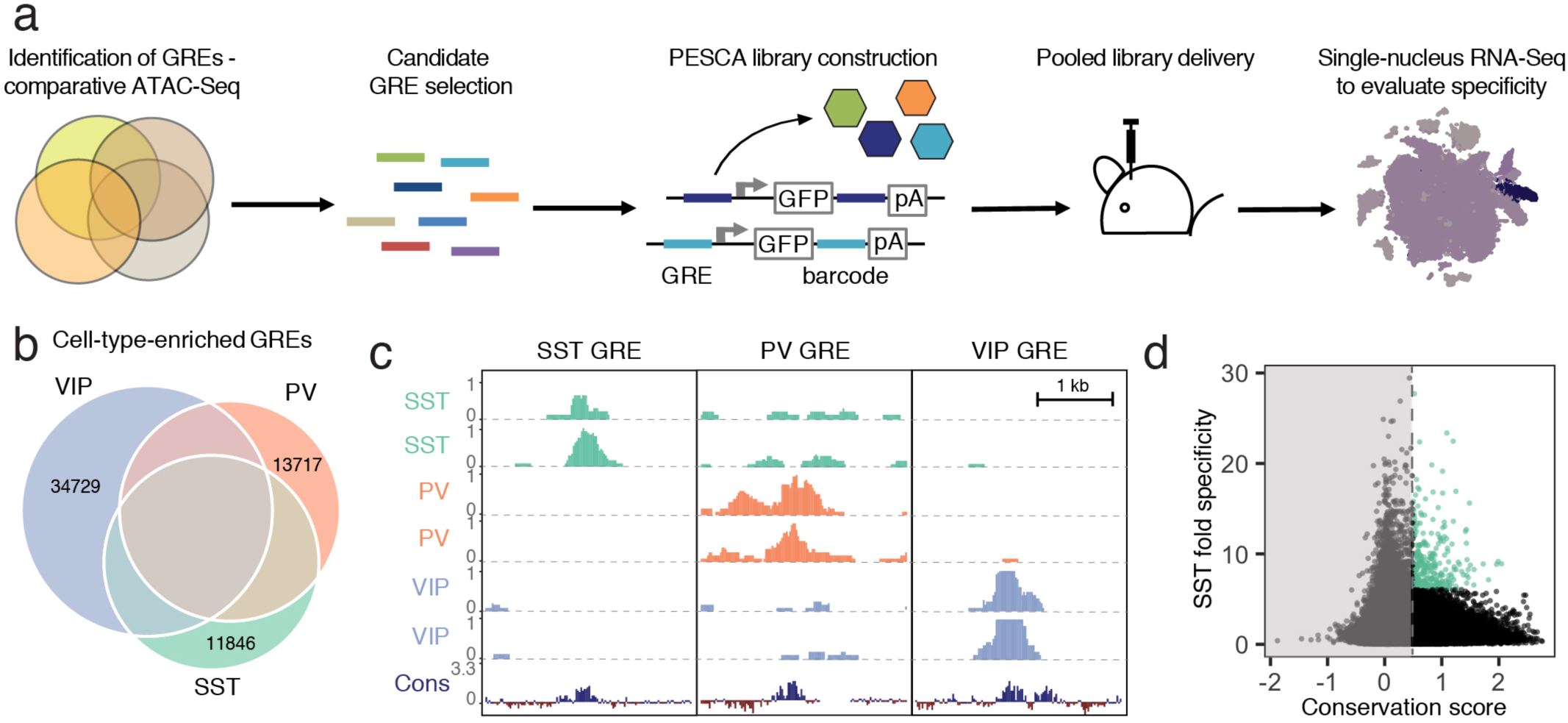
Experimental strategy and GRE selection. **(a)** Paralleled Enhancer Single Cell Assay (PESCA). Comparative ATAC-Seq is used to identify candidate GREs. A library of gene regulatory elements (GREs) is inserted upstream of a minimal promoter-driven GFP. The viral barcode sequence is inserted in the 3’UTR, and the vector packaged into rAAVs. Following *en masse* injection of the rAAV library, the specificity of the constituent GREs for various cell types *in vivo* is determined by single-nucleus RNA sequencing, measuring expression of the barcoded transcripts in tens of thousands of individual cells in the target tissue. Finally, bioinformatic analysis determines the most cell-type-specific barcode-associated rAAV-GRE-GFP constructs. pA = polyA tail. **(b)** Area-proportional Venn diagram of the number of putative GREs identified by ATAC-Seq of purified PV, SST, and VIP nuclei. Overlapping areas indicate shared putative GREs. Non-overlapping areas represent GREs that are unique to a single cell type. **(c)** Representative ATAC-seq genome browser traces of a putative GRE enriched in SST-, PV-, or VIP-interneurons (normalized counts per location). Sequence conservation across the Placental mammalian clade is also shown. **(d)** Putative GREs (n = 323,369) are plotted based on average sequence conservation (phyloP, 60 placental mammals) and SST-specificity (ratio of the average ATAC-Seq signal intensity between SST samples and non-SST samples). Dashed vertical line indicates the minimal conservation value cutoff (0.5). Green coloring indicates the 287 most SST-specific GREs selected for PESCA screening.

We validated the efficacy of PESCA in the murine primary visual cortex by identifying GREs that confine AAV expression to somatostatin (SST)-expressing interneurons and showed that these vectors can be used to modulate neuronal activity selectively in SST neurons. We chose to focus on SST neurons in the brain because this population is known to be diverse and to be composed of several relatively rare subpopulations (*2, 15, 25*), and thus might serve as a good test case. As described below, our findings highlight the utility of PESCA for identifying viral constructs that drive gene expression selectively in a subset of neurons and establish PESCA as a platform of broad interest to the research and gene therapy community, potentially enabling the generation of cell-type-specific AAVs for virtually any cell type.

## Results

### GRE selection and library construction

To identify candidate SST interneuron-restricted gene regulatory elements (GREs), we carried out comparative epigenetic profiling of the three largest classes of cortical interneurons: somatostatin (SST)-, vasoactive intestinal polypeptide (VIP)- and parvalbumin (PV)-expressing cells. To this end, we employed the recently developed isolation of nuclei tagged in specific cell types (INTACT) (*5*) method to isolate purified chromatin from of each of these cell types from the cerebral cortex of adult (6-10-week-old) mice. The assay for transposase-accessible chromatin using sequencing (ATAC-Seq) (*7*), which identifies nucleosome-depleted gene regulatory regions, was then used to identify genomic regions with enhanced accessibility (i.e. peaks) in the SST (n = 57,932), PV (n = 61,108), and VIP (n = 79,124) chromatin samples (**Fig. 1b, c, Supplementary Fig 1, Methods)**. These datasets can be used as a resource to identify putative gene regulatory elements as candidates for driving cell-type-specific gene expression for the numerous subtypes of SST, PV or VIP-expressing intraneurons across diverse cortical regions.

To enrich for GREs that could be useful reagents to study and manipulate interneurons across mammalian species including humans, we started with an expanded list of 323,369 genomic coordinates (**Supplementary Table 1**) representing a union of cortical neuron ATAC-seq accessible regions identified across dozens of experiments in our laboratory (**Methods**, manuscript in preparation). We first filtered this initial set of 323,369 genomic coordinates to exclude GREs with poor mammalian sequence conservation (**Methods, Supplementary Table 1, Supplementary Fig. 2**). The remaining 36,215 genomic regions were ranked by an enrichment of ATAC-seq signal in the SST samples over PV/VIP (**Methods**), and the top 287 most enriched GREs were selected for functional screening to identify enhancers that drive gene expression selectively in SST interneurons of the primary visual cortex (**Fig. 1d, Supplementary Table 2**).

A PCR-based strategy was used to simultaneously amplify and barcode each GRE from mouse genomic DNA (**Methods**). To minimize sequencing bias due to the choice of barcode sequence, each GRE was paired with three unique barcode sequences. The resulting library of 861 GRE-barcode pairs was pooled and cloned into an AAV-based expression vector, with the GRE element inserted 5’ to a promoter driving a GFP expression cassette and the GRE-paired barcode sequences inserted into the 3’ untranslated region (UTR) of the GRE-driven transcript (**Methods, Fig. 2a, Supplementary Fig. 3)**. This configuration was chosen to maximize the retrieval of the barcode sequence during single-cell RNA sequencing, which primarily captures the 3’ end of transcripts. The human beta-globin promoter was chosen since it has previously been used in conjunction with an enhancer to drive strong and specific expression in cortical interneurons (*8*), although the modular cloning strategy is compatible with the use of other promoters. The library was packaged into AAV9, which exhibits broad neural tropism and has previously been used to drive payload expression in cortical neurons (*28*). The complexity of the resulting rAAV-GRE library was then confirmed by next generation sequencing, detecting 802 of the 861 barcodes (93.2%), corresponding to 285 of the 287 GREs (99.3%) (**Fig. 2b**).

**Figure 2.**
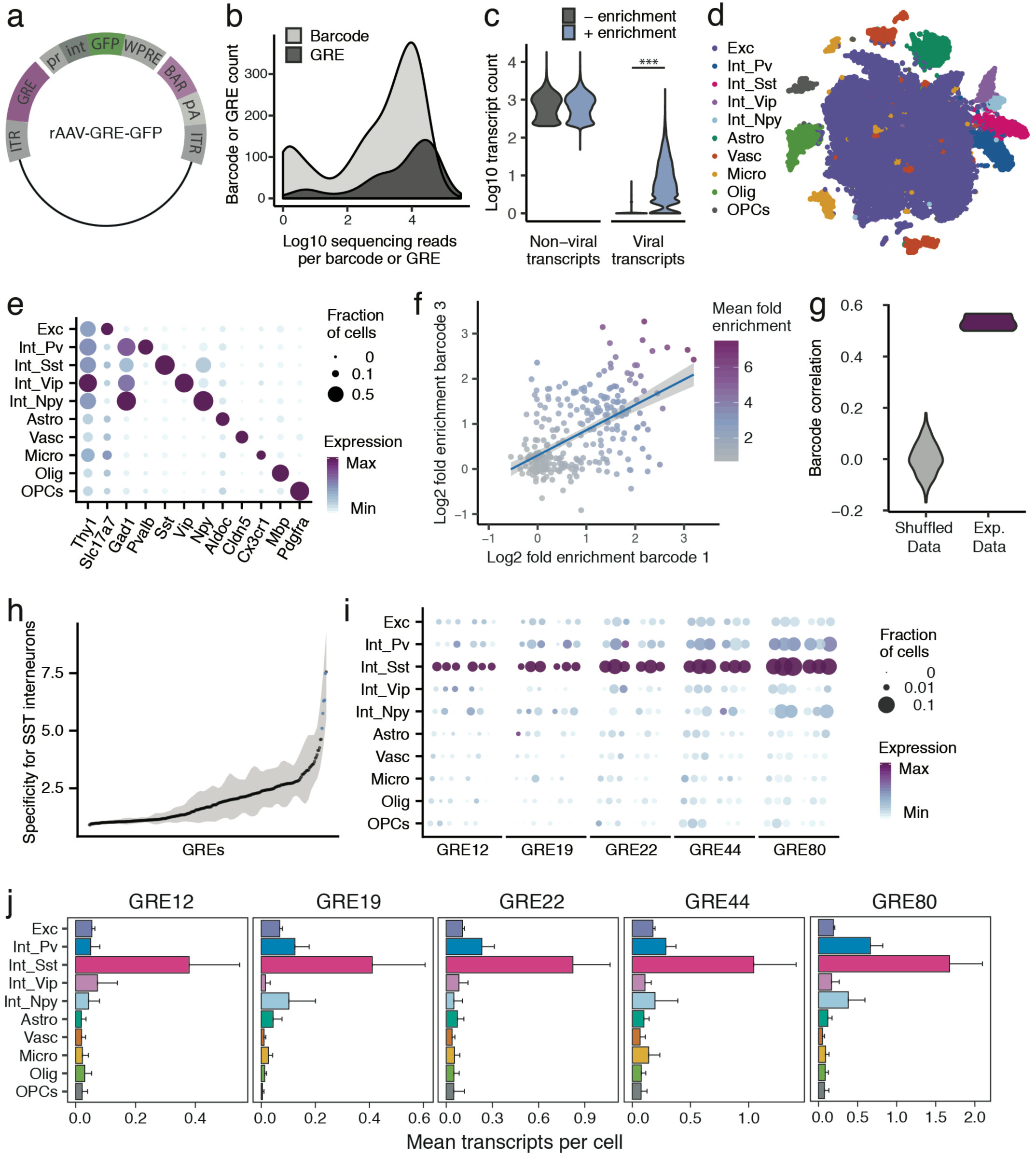
PESCA screen identifies GREs highly enriched for SST^+^ interneurons. **(a)** PESCA library plasmid map. ITR, inverted terminal repeats; GRE, gene regulatory element; pr, HBB minimal promoter; int, intron; GFP, green fluorescent protein; WPRE, Woodchuck Hepatitis Virus post-transcriptional regulatory element; BAR, 10-mer sequence barcode associated with each GRE; pA, polyadenylation signal. **(b)** Library complexity plotted as distribution of the abundance of the 861 barcodes and 287 GREs in the AAV library. Barcodes and GREs were binned by number of sequencing reads attributed to each barcode or GRE within the library. **(c)** Transcript count per nucleus (n = 32,335 nuclei). Sequencing libraries were prepared with or without PCR-enrichment for viral transcripts. PCR enrichment resulted in a 382-fold increase in the number of recovered viral transcripts (p = 0, Mann-Whitney U-test, two-sided) to an average of 15.6 unique viral transcripts per nucleus. Displayed as Log10(Count+1). **(d)** t-SNE plot of 32,335 nuclei from V1 cortex of two animals. Colors denote main cell types: Exc (Excitatory neurons), Pv (PV Interneurons), Sst (SST Interneurons), Vip (VIP interneurons), Npy (NPY Interneurons), Astro (Astrocytes), Vasc (Vascular-associated cells), Micro (Microglia), Olig (Oligodendrocytes), OPCs (Oligodendrocyte precursor cells). **(e)** Marker gene expression across cell types. Color denotes mean expression across all nuclei normalized to the highest mean across cell types. Size represents the fraction of nuclei in which the marker gene was detected. **(f)** Dot plot with each dot representing one GRE (n = 287). The values on each axis represent the Log2 SST fold-enrichment calculated for each GRE based on two of the three barcodes paired with that GRE - barcode 1 on the x-axis, and barcode 3 on the y-axis. Blue line indicates linear fit with 95% confidence intervals (shaded) (r = 0.55, p < 2.2×10^−16^, Pearson’s correlation). Color gradient indicates the average enrichment between the two barcodes. **(g)** Pairwise Pearson correlation between the enrichment values calculated from three sets of barcodes associated with 287 GREs for experimental data (Exp. Data, r = 0.52 ± 0.05, p < 2.2×10^−16^, Pearson’s correlation) and after random shuffling of enrichment values (Shuffled Data, r = 0 ± 0.06). **(h)** GREs ranked by average barcode expression specificity for SST interneurons across three barcodes. Shading indicates the minimal and maximal specificity calculated by analyzing each of the three barcodes associated with a GRE. Blue indicates the five top hits that also passed a statistical test for SST interneuron enrichment (FDR-corrected q < 0.01). **(i)** Expression of the top five hits: GRE12, GRE19, GRE22, GRE44, GRE80. For each GRE, expression values are split into two animals, and, for each animal, into the three barcodes associated with that GRE. Color denotes mean expression across all nuclei normalized to the highest mean across cell types. Size represents the fraction of nuclei in which the marker gene was detected. **(j)** Mean expression of GRE12, GRE19, GRE22, GRE44, and GRE80 across cell types. Error bars, s.e.m.

### PESCA screen identifies GREs highly enriched for SST^+^ interneurons

To quantify the expression of each rAAV-GRE vector across the full complement of cell types in the mouse visual cortex, we used a modified single-nucleus RNA-Seq (snRNA-Seq) protocol to first determine the cellular identity of each nucleus and then quantify the abundance of the GRE-paired barcodes in the transcriptome of nuclei assigned to each cell type. Two adjacent injections (800 nL each) of the pooled AAV library (1 × 10^13^ viral genomes/mL) were first administered to the primary visual cortex (V1) of two 6-week-old C57BL/6 mice. Twelve days following injection, the injected cortical regions were dissected and processed to generate a suspension of nuclei for snRNA-Seq using the inDrops platform (*23, 29*) (**Methods**). A total of 32,335 nuclei were subsequently analyzed across the two animals, recovering an average of 866 unique non-viral transcripts per nucleus, representing 610 unique genes (**Supplementary Fig. 4a, b**).

Since droplet-based high-throughput snRNA-Seq samples the nuclear transcriptome with low sensitivity (*23*), viral-derived transcripts were initially detected in only 3.9% of sampled nuclei. We therefore designed a modified PCR-based approach to enrich for barcode-containing viral transcripts, which yielded deep coverage of AAV-derived transcripts with simultaneous shallow coverage of the non-viral transcriptome. PCR enrichment increased the viral transcript recovery 382-fold in the sampled nuclei, to an average of 15.6 unique viral transcripts, 6.0 unique GRE-barcodes, and 5.7 unique GREs per cell **(Fig. 2b, Supplementary Fig. 4c)**. Using this modified protocol, viral transcripts were identified across 86% of cells (**Supplementary Fig. 4d**), with a high correlation (r = 0.9, p < 2.2×10^−16^) observed between the abundance of each barcoded AAV in the library and the number of cells infected by that AAV (**Supplementary Fig. 4f**), suggesting that GRE sequences did not alter viral tropism and that GRE-driven vectors had broadly similar levels of expression. Only 0.3 ± .06% (mean, stdev) of viral reads did not correspond to any of the known barcodes or could not be uniquely assigned to a barcode (within 2 mismatches), suggesting that this amplification strategy did not grossly change the composition of the viral library.

Nuclei were classified into ten cell types using graph-based clustering and expression of known marker genes **(Methods, Fig. 2c, d, Supplementary Fig. 5**). The average expression of each viral-derived barcoded transcript was analyzed across all ten cell types, and an enrichment score was calculated from the ratio of expression in *Sst*^*+*^ nuclei compared to all *Sst* ^*-*^ nuclei. As expected, sets of three barcodes associated with the same GRE showed highly statistically correlated enrichment scores (r = 0.52 ± 0.05, p < 2.2×10^−16^) (**Fig. 2e, f, Supplementary Fig. 6**), which were significantly lower when barcodes were randomly shuffled (shuffled r = 0.002 ± 0.06; Wilcox test between data and shuffled data, p = 0.003).

Having confirmed a robust, non-random correlation in enrichment scores among the three barcodes associated with each GRE, we next computed a single expression value for each of the 287 viral drivers by aggregating expression data from three barcodes associated with the same GRE, and carried out differential gene expression analysis between *Sst*^*+*^ and *Sst*^*-*^ cells for each rAAV-GRE. Differential gene expression analysis between *Sst*^*+*^ and *Sst* ^*-*^ cells for each rAAV-GRE revealed a marked overall enrichment of viral-derived transcripts in the *Sst*^*+*^ subpopulation (**Supplementary Fig. 7a**). As expected, a high correlation was observed between GRE-specific enrichment scores across two animals (r = 0.54, p < 2.2×10^−16^) (**Supplementary Fig. 7b**). Among the 287 GREs tested, several viral drivers were identified that promoted highly specific reporter expression in the *Sst*^*+*^ subpopulation (q < 0.01, fold-change > 7, **Fig. 2h-j, Supplementary Fig. 7c-e**). To assess how the abundance of each GRE in the library impacts our ability to detect cell-type-specific expression, we analyzed the specificity of each GRE as a function of the number of transcripts retrieved. We observed that highly abundant GRE-driven transcripts were more likely to be significantly enriched in SST^+^ cells, suggesting that we may not have had sufficient power to assess the cell-type-specificity of the less abundant GREs in the library (**Supplementary Fig. 7f**). Consistent with this observation, computationally subsampling the number viral transcripts across our most cell-type-specific GREs gradually reduced our ability to statistically detect their enrichment in *Sst*^*+*^ cells (**Supplementary Fig. 8)**. These observations suggest that the expression of sparsely detected GRE-driven transcripts may not be sufficient to allow evaluation of cell-type-specificity and that by increasing sequencing depth we may be able to screen and evaluate a larger number of GREs.

### *In situ* characterization of rAAV-GRE reporter expression

We next sought to validate the cell-type-specificity of the resulting hits using methods that do not rely on single-cell sequencing-based approaches. To this end, we selected three of the top five viral drivers (GRE12, GRE22, GRE44), as well as a control viral construct lacking the GRE element (ΔGRE), for injection into V1 of adult transgenic Sst-Cre; Ai14 mice, in which SST^+^ cells express the red fluorescent marker tdTomato (**Supplementary Table 3**).

Fluorescence analysis twelve days following injection with rAAV-[GRE12, GRE22 or GRE44]-GFP revealed strong yet sparse GFP labeling centered around cortical layers IV and V (**Fig. 3a-c).** By contrast, the control rAAV-ΔGRE-GFP showed a strikingly different pattern of GFP expression concentrated around the sites of injection, with expression in a larger number of cells (**Fig. 3d**). Many rAAV-GRE12/22/44-GFP virally infected cells were SST-positive, as indicated by the high degree of overlapping GFP and tdTomato expression: 90.7% ± 2.1% for rAAV-GRE12-GFP (170 cells, 4 animals); 72.9 ± 4.2% for rAAV-GRE22-GFP (1164 cells, 3 animals), and 95.8 ± 0.6% for rAAV-GRE44-GFP (759 cells, 4 animals). (**Fig. 3e, f, Supplementary Fig. 9**). By contrast, we observed that 27.2 ± 1.9% of GFP^+^ cells also expressed tdTomato following rAAV-ΔGRE-GFP infection (2066 cells, 3 animals, **Fig. 3e, f**). Although the 27.2% overlap between rAAV-ΔGRE-GFP expression and SST^+^ cells suggests that our vector has some baseline preference for SST^+^ interneurons, the insertion of GRE12, GRE22 and GRE44 serves to effectively restrict AAV payload expression to SST^+^ interneurons. To show that not all the enhancers exhibited strong SST-specificity using our viral vector, we cloned the mDlx5/6 enhancer whose expression was restricted to a broader population of inhibitory neurons (*8*). We injected the rAAV2/9-mDlx5/6-GFP vector and observed that 42.9 % of GFP^+^ cells were also positive for tdTomato (1977 cells, 3 animals, **Supplementary Fig. 10**). It is notable that the GREs seemingly not only promote expression in SST^+^ cells but also greatly reduce background expression in SST negative cells, indicating both enhancer and repressor functionality. Consistent with this hypothesis, the incorporation of GRE12, GRE22 and GRE44 into the rAAV both increased the number of SST^+^ GFP^+^ cells (1.7-2-fold) and dramatically (3-32-fold) decreased the number of SST^-^ cells that expressed GFP (**Fig. 3g, Supplementary Fig. 11**). To further investigate the specificity of our viral drivers among cortical interneuron cell types we injected each construct into Vip-Cre; Ai14^+^ mice in which all VIP^+^ cells express tdTomato, or used fluorescence antibody staining to label PV-expressing cells (**Supplementary Fig. 12)**. Fluorescent signal analysis indicated the percentage of GFP^+^ cells that were either VIP^+^ or PV^+^ (rAAV-SST12-GFP^+^ [2.6 ± 2.6%], rAAV-GRE22-GFP^+^ [3.5 ± 2.0%] and rAAV-GRE44-GFP^+^ [6.0 ± 2.7%], **Fig. 3h**). These findings confirm that among major interneuron cell classes, all three GRE-driven vectors are highly SST-specific.

Because at least five subtypes of cortical SST^+^ interneurons have previously been identified based on the laminar distribution of their cell bodies and projections (*15, 30*), we investigated the laminar distribution of GFP-expressing cells for the three Sst-enriched viral drivers. Intriguingly, the majority of rAAV-GRE12-GFP^+^ and rAAV-GRE44-GFP^+^ SST^+^ cells were found to reside in layers IV and V, which was distinct from the distribution observed for the full SST^+^ cell population in visual cortex (p = 1.3×10^−6^, p < 2.2×10^−16^, respectively, Mann-Whitney *U* test, two-tailed, **Fig. 3i, Supplementary Fig. 13**). By contrast, rAAV-ΔGRE-GFP labeled SST^+^ cells as well as other neuronal subtypes across all layers, suggesting that the confined labeling of rAAV-GRE12-GFP and rAAV-GRE44-GFP to layer IV and V was likely due to restricted gene expression and not restricted viral tropism. These findings suggest that PESCA may support the isolation of viral drivers capable of discriminating between fine-grained cell-types within a given interneuron cell class.

**Figure 3.**
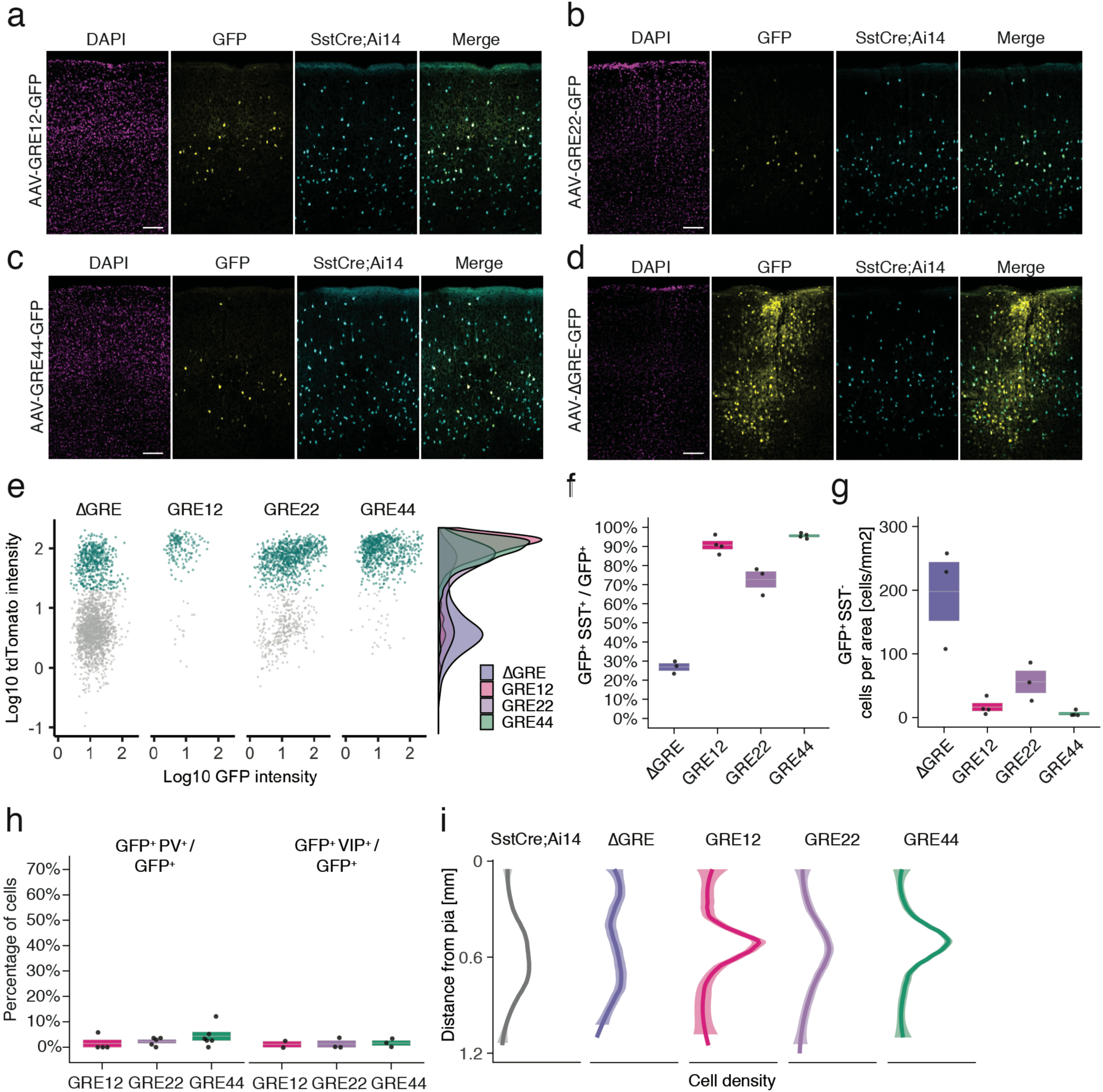
*In situ* characterization of rAAV-GRE reporter expression. (**a-d**) Fluorescent images from adult Sst-Cre; Ai14 mouse visual cortex twelve days following injection with rAAV-GRE-GFP as indicated. Scale bars 100 µm. **(e)** Identification of rAAV-GRE-GFP^+^ cells that express tdTomato (SST^+^). Each dot represents a GFP^+^ cell (n = 2066, 172, 1164, and 765, for AAV-[ΔGRE, GRE12, GRE22, GRE44]-GFP, respectively). Cyan indicates tdTomato^+^ (SST^+^) cells. Distribution of cell frequency across tdTomato intensity is plotted on the right for each construct. **(f)** Quantification of the fraction of GFP^+^ cells that are SST^+^. Each dot represents one animal. Box plot represents mean ± standard error of the mean (s.e.m). Values are 27.2 ± 1.9%, 90.7 ± 2.1, 72.9 ± 4.2%, and 95.8 ± 0.6% for AAV-[ΔGRE, GRE12, GRE22, GRE44]-GFP, respectively. **(g)** Quantification of the number of GFP^+^ SST^-^ cells normalized for area of infection. Each dot represents one animal. Box plot represents mean ± standard error of the mean (s.e.m). Values are 198.0 ± 46.0, 16.4 ± 6.2, 56.0 ± 17.3 and 6.1± 2.1 cells/mm^2^ for AAV-[ΔGRE, GRE12, GRE22, GRE44]-GFP, respectively. **(h)** Quantification of the fraction of GFP^+^ cells that are PV^+^ or VIP^+^. Box plot represents mean ± standard error of the mean (s.e.m). Fraction of AAV-GRE-GFP^+^ cells that are PV^+^ is 1.4 ± 1.4%, 2.2 ± 0.7, and 4.3 ± 1.7% for AAV-[GRE12, GRE22, GRE44]-GFP, respectively. Similarly, the fraction of AAV-GRE-GFP^+^ cells that are VIP^+^ is 1.2 ± 1.2%, 1.3 ± 1.3%, and 1.7 ± 1.0% for AAV-[GRE12, GRE22, GRE44]-GFP^+^ cells, respectively. **(i)**Distribution of the location of GFP-expressing cells as function of distance from the pia. Gray represents SST^+^ cells (n = 2648); Colored lines represents GFP^+^ SST^+^ cells (n = 2066, 172, 1164, and 765, respectively, for AAV-[ΔGRE, GRE12, GRE22, GRE44]-GFP). Shading represents the 95% confidence interval.

### Electrophysiological characterization of rAAV-GRE-GFP-expressing SST subtypes

In addition to variability in laminar distribution, different electrophysiological phenotypes have also been observed in cortical SST interneurons (*31, 32*). To determine whether AAV-GRE reporters can be used to distinguish electrophysiologically distinct SST subtypes, we injected our most cell-type-restricted construct rAAV-GRE44-GFP into the visual cortex of adult Sst-Cre; Ai14 mice and obtained whole-cell current-clamp recordings from double GFP- and tdTomato-positive neurons (rAAV-GRE44-GFP^+^), as well as immediately nearby tdTomato-positive but GFP-negative cells (rAAV-GRE44-GFP^-^).

Our recordings indicate that both rAAV-GRE44-GFP^+^ and rAAV-GRE44-GFP^-^ SST^+^ neurons display the properties of adapting SST interneurons with high input resistances and features consistent with those previously reported for deep layer cortical SST neurons(*31, 33*) (**Fig. 4a, b**). However, rAAV-GRE44-GFP^+^ SST neurons are distinct with respect to several electrophysiological parameters. The action potentials of rAAV-GRE44-GFP^+^ SST neurons are significantly broader than those of rAAV-GRE44-GFP^-^ SST neurons (**Fig. 4c, d**), perhaps due to differences in expression of specific channels in these subgroups of SST neurons, such as voltage-activated potassium channels, and BK calcium-activated potassium channels(*34, 35*). Furthermore, rAAV-GRE44-GFP^+^ SST neurons have a lower rheobase, and fire action potentials with a slower rising phase, and at lower maximal frequencies compared to rAAV-GRE44-GFP^-^ SST neurons **(Fig. 4a, d, Supplementary Table 4)**. Although we cannot confirm that GRE44 expression is restricted to a specific transcriptionally defined subtype of SST interneurons, our electrophysiology experiments further emphasize the potential of PESCA to target functionally distinct subgroups of previously defined interneuron types.

**Figure 4.**
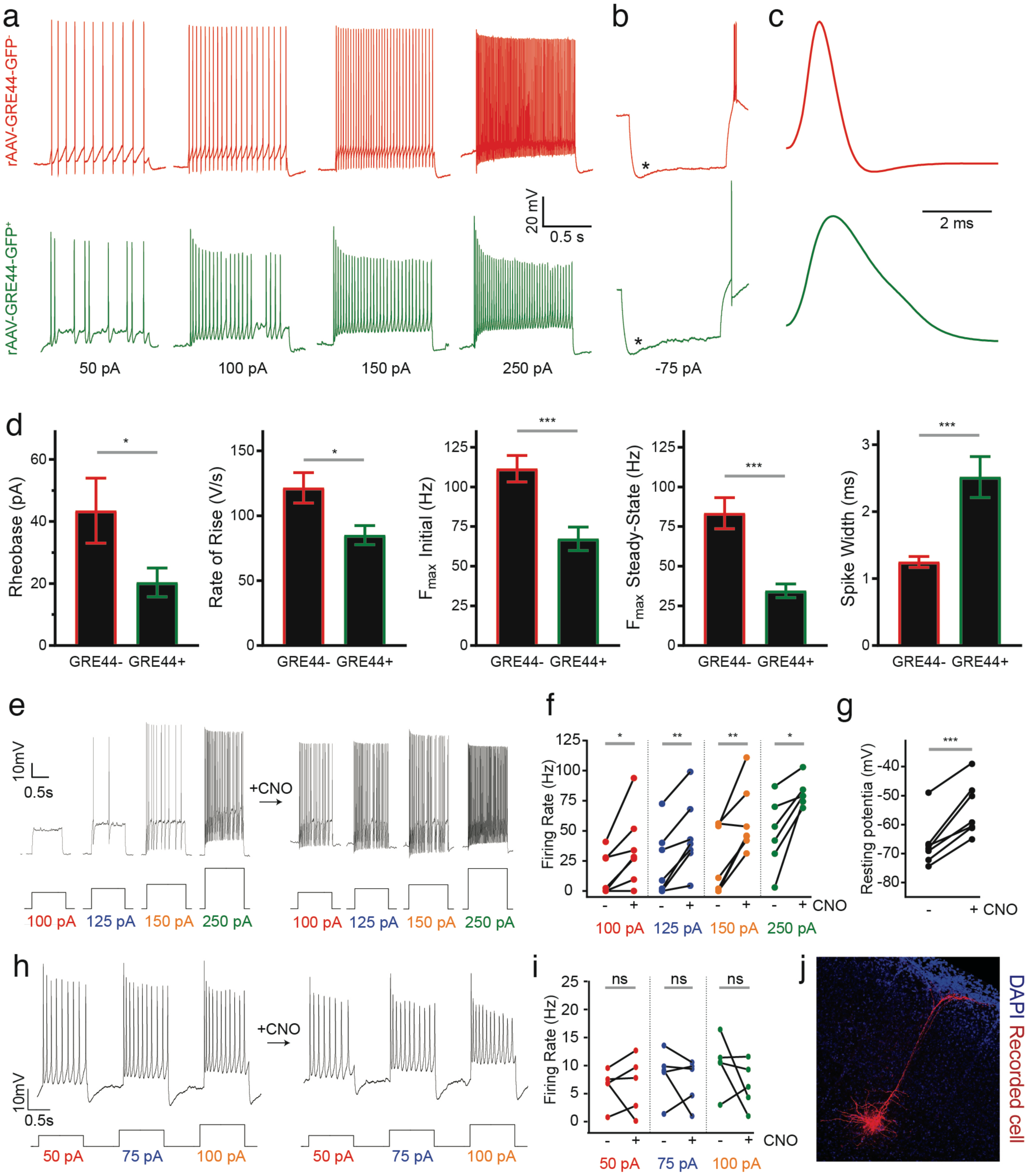
Electrophysiology of neurons expressing an rAAV-GRE-driven reporter and modulation of neuronal activity with rAAV-GREs. **(a)** Representative current-clamp recordings from SST neurons in the visual cortex of Sst-Cre; Ai14 mice injected with rAAV-GRE44-GFP. *Top*: Representative traces from a cortical SST neuron with Cre-dependent expression of red fluorescent protein (RFP), in response to 1000 ms depolarizing current injections as indicated in black (“GRE44-”). *Bottom*: Traces from a RFP^+^ SST neuron with GRE44-driven expression of GFP (“GRE44^+^”). GRE44-SST neurons were only recorded in the immediate vicinity of GRE44^+^ SST neurons. **(b)** Recordings from GRE44^+^ and GRE44^-^ neurons in response to hyperpolarizing, 1000 ms currents. Asterisks indicate the sag likely due to the hyperpolarization-activated current *I*_h_. Rebound action potentials following recovery from hyperpolarization, likely due to low-threshold calcium spikes mediated by T-type calcium channels, were also present in cells of both groups. Same scale as (a). **(c)** Broader action potentials in GRE44^+^ SST neurons (bottom) compared to GRE44^-^ SST neurons (top). Same vertical scale as (a) and (b). **(d)** Electrophysiological properties that differ between GRE44^+^ (n=16 cells from 5 mice) and GRE44^-^ (n=16 cells from 4 mice) SST neurons, including rheobase (minimal amount of current necessary to elicit a spike), maximal rate of rise during the depolarizing phase of the action potential, the initial and steady state firing frequencies (both measured at the maximal current step before spike inactivation), and spike width (measured as the width at half-maximal spike amplitude). * p < 0.05; *** p < 0.001, paired t-test, two-tailed. **(e)** Representative current-clamp recordings from AAV-GRE12-Gq-tdTomato^+^ cells before and during CNO application. **(f)** Increased firing rates of AAV-GRE12-Gq-tdTomato^+^ cells evoked by depolarizing current injections upon bath application of CNO (3 animals, 6-7 cells). * p < 0.05; ** p < 0.01, paired t-test, two-tailed. **(g)** Robust depolarization of AAV-GRE12-Gq-tdTomato^+^ cells upon bath application of CNO (3 animals, 6-7 cells). *** p < 0.001, paired t-test, two-tailed. **(h)** Representative recordings from nearby uninfected pyramidal neurons in the visual cortex of mice that were injected with AAV-GRE-12-Gq-tdTomato^+^, before (top) and during CNO application (bottom). **(i)** Firing rates of pyramidal neurons during CNO application remain unchanged (3 animals, 5 cells). ns, p > 0.05, paired t-test, two-tailed. **(j)** Representative image of nearby uninfected pyramidal neuron that was recorded from

### Modulation of neuronal activity with rAAV-GREs

Finally, we evaluated whether the identified SST^+^ neuron-restricted viral drivers support sufficiently high and persistent levels of payload expression to effectively modulate SST^+^ cell physiology. Designer receptors exclusively activated by designer drugs (DREADDs) are a commonly employed viral payload used to dynamically regulate neuronal activity in response to the synthetic ligand clozapine-N-oxide (CNO)(*36*). We therefore injected the visual cortex of adult wild-type mice (6-8-week-old) with rAAV-GRE12-Gq-DREADD-tdTomato, a construct in which GRE12 drives the expression of an activating DREADD as well as tdTomato. GRE12 was chosen for this assay as it drives the weakest expression of the three evaluated GREs (**Fig. 2e)** and thus, if it effectively drives DREADD expression, the other GREs might be expected to as well. We obtained electrophysiological recordings from tdTomato^+^ cells of acute cortical slices in a whole-cell, current-clamp configuration two weeks post-injection. All tdTomato^+^ cells showed striking sensitivity to CNO, as indicated by significantly increased firing rates in response to depolarizing current steps and depolarized resting membrane potentials (**Fig. 4e-g**). To ensure that increases in firing rate upon CNO application were specific to infected SST^+^ neurons, we obtained recordings from nearby uninfected pyramidal neurons that were identified by morphology and found that there was no statistically significant increase in firing rate upon CNO application (**Fig. 4h-j**). These data demonstrate the ability of GRE-driven SST^+^ neuron-specific reagents to robustly and specifically modulate the activity of SST^+^ cells in non-transgenic animals.

## Discussion

The PESCA platform extends previous paralleled reporter assays (*17–22*) carried out using bulk tissue or sorted cells by including a single-cell RNA-seq-based readout (*1*–*4, 23*–*27*) to evaluate the cell-type-specificity of gene expression. This represents a significant advancement over current approaches to viral vector design, as it enables the rapid *in vivo* screening of hundreds of GREs for enhanced cell-type-specificity without needing transgenic tools to evaluate their specificity. In this study, we applied PESCA to identify enhancer elements that robustly and specifically drive gene expression in a rare SST^+^ population of GABAergic interneurons in the mouse central nervous system, although further work is needed to identify which specific molecular subtypes of SST interneurons are targeted. Since the vectors used in this PESCA screen in the absence of GREs show broad expression in the murine V1, the GREs we identified likely function to both enhance and restrict viral expression by a mechanism that remains to be explored.

In the future, several factors should be considered to facilitate the further optimization of the PESCA methodology for the development of cell-type-specific vectors. The selection of candidate GREs for screening will benefit from the systematic profiling of additional cell types by traditional or single-cell ATAC-Seq methods. In this regard, consideration of a published ATAC-Seq dataset from excitatory neurons(*5*) could have served to refine our starting GRE set by excluding approximately half of the screened GREs from our initial pool. This is particularly relevant insofar as the ability to assess the GRE library depends on the number of cells sequenced from the target and non-target populations and the sequencing depth, as the coverage of each GRE will be inversely proportional to the number of GREs screened. In the screen described here, we estimate having sufficient power to assess approximately 2/3 of the 287 GREs at the reported sequencing depth (**Supplementary Fig. 7**).

If a robust method of specifically isolating RNA from the target cell population is available, screening the PESCA library by sequencing pooled RNA from all target versus all non-target cells would provide less expensive and potentially more scalable approach. However, by averaging across multiple non-target cell types, such an approach could be confounded by the presence of rare, highly expressing non-target cells. Similarly, among single-cell methods, single-nucleus RNA-seq is technically simpler and can be used on frozen tissue but recovers fewer transcripts per cell compared to single-cell RNA-seq.

Finally, once candidate PESCA hits have been identified, we suggest evaluating several in follow-up assays at multiple titers to identify which among these hits have the desired intensity and specificity of protein expression. In this regard, the snRNA-seq PESCA screen identified GRE12, GRE22 and GRE44 as, respectively, 8.3-, 9.1- and 7.2-fold more highly expressed in SST^+^ compared to SST^-^ cells, whereas these GREs showed distinct specificity for SST^+^ cells (91%, 73% and 96% respectively, **Fig. 3f**) when evaluated at the protein level, a finding which could be attributed to a variety of factors.

Given current evidence that the mechanisms of gene regulatory element function are conserved across tissues and species, it is likely that PESCA can be readily applied to other neuronal or non-neuronal cells types, diverse model organisms, tissues, and viral types. Moreover, PESCA is not limited to GRE screening; the method can be easily adapted to assess the cell-type-specificity of viral capsid variants or other mutable aspects of viral design. Indeed, the PESCA library cloning strategy is largely vector- and capsid-independent, allowing for the use of different promoters or serotypes. Our choice of capsid and promoter was driven by previous work using AAV9 and the minimal beta-globin promoter to drive expression in cortical interneurons (*8*), however, different capsids or promoter may be preferred for targeting other cell types.

In conclusion, our study addresses the urgent practical need for new tools to access, study, and manipulate specific cell types across complex tissues, organ systems, and animal models by providing a screening platform that can be used to rapidly supply such tools as needed. Moreover, as the promise of gene therapy to treat and cure a broad range of diseases is being realized, PESCA has the potential to pave the way for a new generation of targeted gene therapy vehicles for diseases with cell-type-specific etiologies, such as congenital blindness, deafness, cystic fibrosis, and spinal muscular atrophy.

## Acknowledgements

We thank D. Tom for help with image analysis, Dr. W. Renthal and members of the Greenberg lab for discussions, the HMS Single Cell Core for single-nucleus RNA-seq sample processing, Boston Children’s Hospital Viral Core for AAV packaging, Dr. B. Bean for feedback on electrophysiology, and the Harvard NeuroDiscovery Center Enhanced Neuroimaging Core for imaging and image analysis. This work was supported by the National Institute of Health BRAIN Initiative grant R01 MH114081-01, NIH grant T32GM007753 to M.A.N. and NIH Training in the Molecular Biology of Neurodegeneration grant 5T32AG000222-23 to S.H.

## Author Contributions

S.H. conceived the study and designed the experiments. H.S. performed ATAC-Seq. M.A.N. aligned and called peaks for ATAC-seq data; S.H. calculated ATAC-seq enrichment and selected elements for screen. S.H. cloned the library and performed the PESCA screen. M.A.N. mapped snRNA-seq data and quantified viral barcodes. S.H. analyzed snRNA-seq data. S.H., C.K. and O.F.W. cloned individual viral constructs. O.F.W. performed stereotactic injections and tissue sectioning. S.H. and E.G.A. performed imaging and image analysis. C.P.T. performed and analyzed electrophysiology recordings. J.G., C.D.G., E.C.G. and M.E.G. advised on the study. S.H., C.P.T., M.A.N, E.C.G. and M.E.G. wrote the manuscript.

